# An atlas of infiltrated B-lymphocytes in breast cancer revealed by paired single-cell RNA-sequencing and antigen receptor profiling

**DOI:** 10.1101/695601

**Authors:** Qingtao Hu, Yu Hong, Pan Qi, Guangqing Lu, Xueying Mai, Sheng Xu, Xiaoying He, Yu Guo, Linlin Gao, Zhiyi Jing, Jiawen Wang, Tao Cai, Yu Zhang

**Author notes:** Correspondence should be addressed to Y. Z. Theses authors contributed equally to this work (Qingtao Hu, Yu Hong, Pan Qi).

## Abstract

While it has been well-recognized that T-cell mediated adaptive cellular immunity plays important roles in cancer immune response and tumor control, the roles of B lymphocytes in tumor development and therapy have only been proposed until recently, and are still mostly controversial. To gain mechanistic insights into the origin and dynamics of tumor infiltrated immune cells, especially B lymphocytes, we combine single-cell RNA-sequencing and antigen receptor lineage analysis to characterize a large number of triple-negative breast cancer (TNBC) infiltrated immune cells and present a comprehensive atlas of infiltrated B-lymphocytes in TNBC, the most aggressive breast cancer subtype. We demonstrate that TNBC infiltrated B cells showed more mature and memory B cell characteristics, as well as high clonality and extensive IgH class switching recombination and somatic hypermutations. The B cell signatures based on single-cell RNA-seq results are significantly associated with improved survival for TNBC patients and provide better prognostication than classic single B cell markers (CD19 or CD20). Further dissection of the mechanisms regulating the functions and dynamic distribution of tumor infiltrated B cell populations will provide new clues for tumor immunotherapy.

## Introduction

Although the roles of immune system in tumor development and therapy have been proposed long time ago, they have only been recognized and mechanistically investigated recently(Finn, 2018; Wei et al., 2018). Both innate immune system (macrophages, neutrophils, master cells, myeloid cells, dendritic cells, and natural killer cells) and adaptive immune system (T and B lymphocytes) could contribute to the establishment of an immunosuppressive tumor microenvironment, which is one of cancer hallmarks (Hanahan and Weinberg, 2011). More specifically, the adaptive immune system creates immunological memory after an initial response to a specific antigen (e.g. tumor antigen), and leads to an enhanced response to subsequent encounters with that antigen to provide long-lasting protection, therefore is supposed to play important roles during tumor development in the cancer immunosurveillance hypothesis and immunoediting hypothesis(Dunn et al., 2004).

While it has been well accepted that T cell mediated adaptive cellular immunity plays important roles in immune response for tumors(Thommen and Schumacher, 2018), the roles of B cells in tumor development and therapy, both positive and negative, have been only proposed recently and are still mostly controversial(Liu et al., 2018; Sarvaria et al., 2017; Shen et al., 2018; Tsou et al., 2016; Wouters and Nelson, 2018). B cells can positively participate in tumor immunology through antibody production, antigen-presentation, cytokine and chemokine production, and immunoregulation mechanisms(Li et al., 2011; Moynihan et al., 2016; Wittrup, 2017). On the other hand, B regulatory (Breg) cells play important roles in maintaining immune homeostasis mainly by secreting cytokines (e.g. IL-10) and/or interacting with target cells(Rosser and Mauri, 2015; Sarvaria et al., 2017; Tsou et al., 2016).

While tumor-infiltrating T cells have been characterized in several human cancer types by single-cell RNA sequencing recently(Azizi et al., 2018; Guo et al., 2018; Li et al., 2018; Sade-Feldman et al., 2018; Savas et al., 2018; Tirosh et al., 2016; Zhang et al., 2018; Zheng et al., 2017a), a comprehensive atlas of tumor-infiltrating B cells is still missing. B cell execrates its functions mainly by recognizing antigen through B-cell receptors (BCRs, antibodies). The diversity of BCRs was generated by V(D)J recombination, somatic hypermutation (SHM), and class switch recombination (CSR) during B cell development and differentiation(Alt et al., 2013). Antigen receptor repertoire analysis provides direct developmental lineage information(Meng et al., 2017). To gain mechanistic insights into the origin and dynamics of infiltrated immune cell, especially B cell, subgroups in TNBC, we combined antigen receptor clonal lineage analysis and single-cell RNA-sequencing analysis to present an atlas of BCR repertoire, clonal lineage, and transcriptional characteristics for TIL-Bs, which will be a foundation for studies of B-cell tumor immunology.

## Results

Human breast cancer is a heterogeneous disease and contains several histologically different subtypes(Polyak, 2011; Tong et al., 2018). We FACS-analyzed infiltrated hCD45^+^ immune cells in fresh isolated breast cancer samples and found that the presence of infiltrated hCD20^+^ B cells was significantly higher in TNBCs than in other breast cancer subtypes (FigS1). To investigate the roles of infiltrated immune cells, especially B cells in human breast cancer, we purified hCD45^+^ cells from surgically isolated breast cancer tissues (BC) and corresponding peripheral blood (PBMC) samples from six TNBC (TNBC1-6 and PBMC1-6), three luminal-A breast cancer (LABC7-9 and PBMC7-9), and one HER2-positive breast cancer (HER2BC10 and PBMC10) treatment-naïve patients (Fig1A), in which significant numbers of infiltrated hCD45^+^ and hCD20^+^ cells could be obtained. Among the samples from these ten patients, nine pairs of them (TNBC2-5, LABC7-9, HER2BC10, and PBMC2-10) were used to prepare 5’ single cell RNA-sequencing libraries by droplet-based (10X Genomics) technology (Zheng et al., 2017b), while one pair (TNBC1 and PBMC1) was used for 3’ library construction (Table S1). In addition, single cell immune repertoire information, including both BCRs (B-cell receptors) and TCRs (T-cell receptors), was also obtained for all 5’ libraries (Table S2).

**Figure 1.**
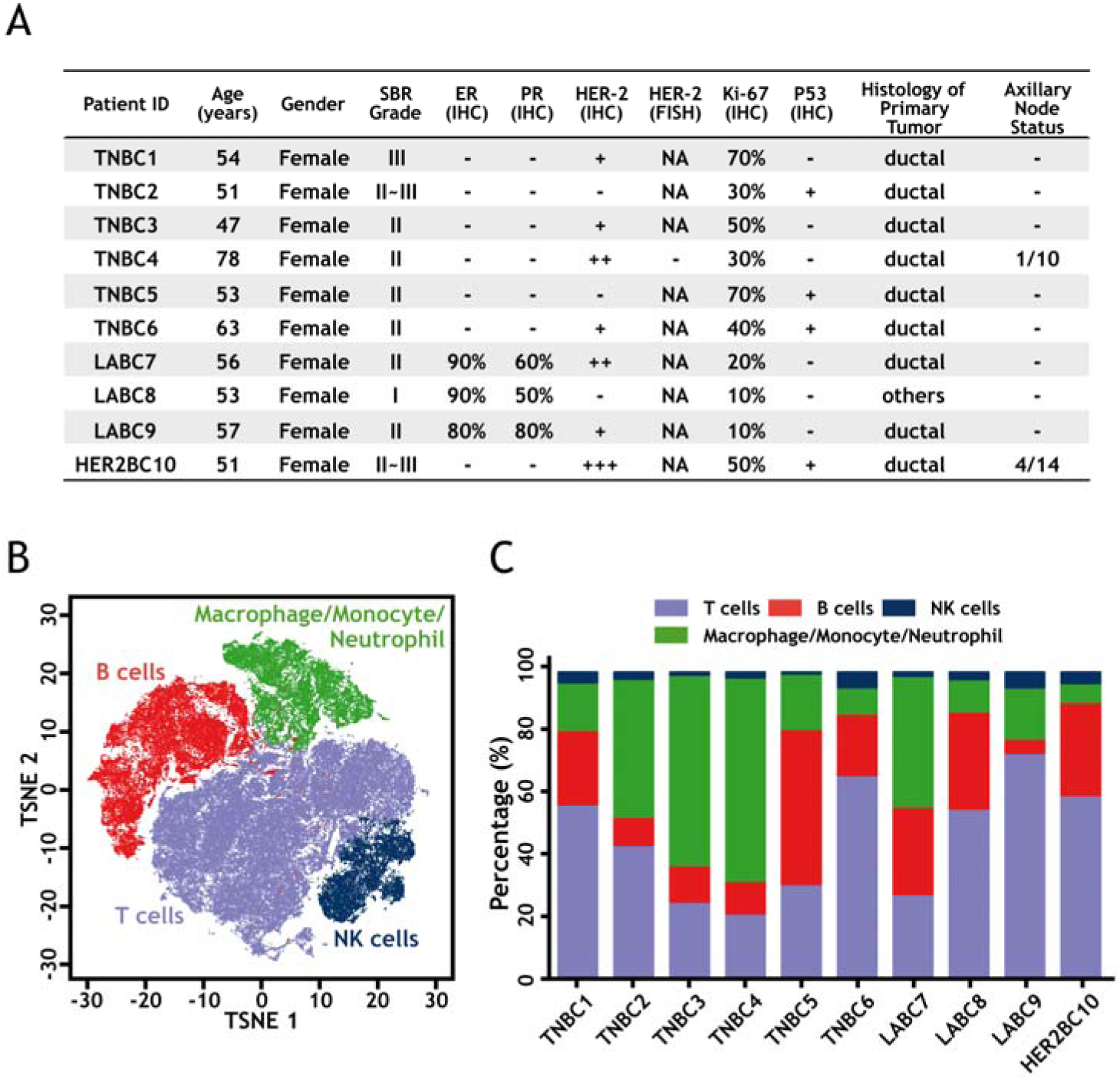
Heterogeneity of infiltrated immune cells in human breast cancers. A: Clinical information of the breast cancer patients involved in current study. B: The t-SNE projection of breast cancer infiltrated immune cell atlas constructed from all patient samples. Each dot represents a cell and major cell types are marked according to FigS3A. C: Distribution of major infiltrated immune cell types in each breast cancer sample.

Histological analysis, immunohistochemistry (IHC), and fluorescence in situ hybridization (FISH) were used to confirm the subtypes of breast cancer samples (Fig1A). Whole exome sequencing (WES) analysis was also performed for both BC and PBMC samples from the six TNBC patients to identify germline and somatic mutations (FigS2 and TableS3).

After quality control and removing potential cell doublets(McGinnis et al., 2019), we obtained totally 44,497 cells with 1,573 median genes per cell from ten tumor samples and 68,441 cells with 1,651 median genes per cell from the corresponding peripheral blood samples (TableS1). Unsupervised clustering(Butler et al., 2018) at low resolution on these cells from all sequenced samples revealed four major cellular clusters including T cells (marked by CD3D expression), B cells (CD20), NK cells (NKG7), and macrophage/monocyte/neutrophil (CD14) (Fig1B and FigS3A). Cells from different samples contributed similarly to each cluster, suggesting lack of sample batch effect (FigS3B and FigS3C). The distributions of different infiltrated lymphocyte clusters in each tumor were heterogeneous across patients (Fig1C). For example, the percentages of B cells in tumor samples varied between 4.6-50.5%.

By the 10x Genomics 5’ V(D)J and gene expression chromium platform, we could identify the rearrangement status of antigen receptor loci in each B and T cell. After quality control, we totally assembled rearranged BCRs and TCRs for 33,509 B cells and 58,095 T cells in all samples except TNBC1/PBMC1, respectively (TableS2). Among them, there are 5,951 and 16,485 B cells containing a single productive IgH allele for BC and PBMC samples, respectively. As TNBC tumors have significantly more infiltrated B cells, in current study, we focused mainly on B cells from the five TNBC patients (TNBC2-6 and PBMC2-6). Analysis of the single productive IgH rearrangement from 3,695 infiltrated B cells in TNBCs and 8,037 B cells from their corresponding PBMCs did not reveal significantly different VH, DH, and JH gene usage between tumor and PBMC samples (FigS4A). However, B cells infiltrated in TNBC contained significantly higher percentage of IgG-positive cells and lower percentage of IgM- and IgD-positive cells than those in peripheral blood (Fig2A and FigS4B). In agree with this, the percentages of B cells have germline productive IgH alleles are significantly lower in TNBC tumors (Fig2A), suggesting the B cells infiltrated in TNBC have encountered antigens and experienced BCR activation. Somatic hypermutation (SHM) analysis(Alamyar et al., 2012; Gupta et al., 2015) demonstrated that overall TNBC infiltrated B cells have significantly more IgH mutations than in PBMC, although the mutation spectrum is similar between TNBC and PBMC B cells (Fig2B and FigS4C-D). Interestingly, such difference is not only due to lower percentages of germline IgH containing cells in tumor infiltrated B cells. The SHM rate in non-germline and non-switched (IgM- and IgD-positive) B cells is also significantly higher in tumor than that in PBMC (Fig2B).

**Figure 2.**
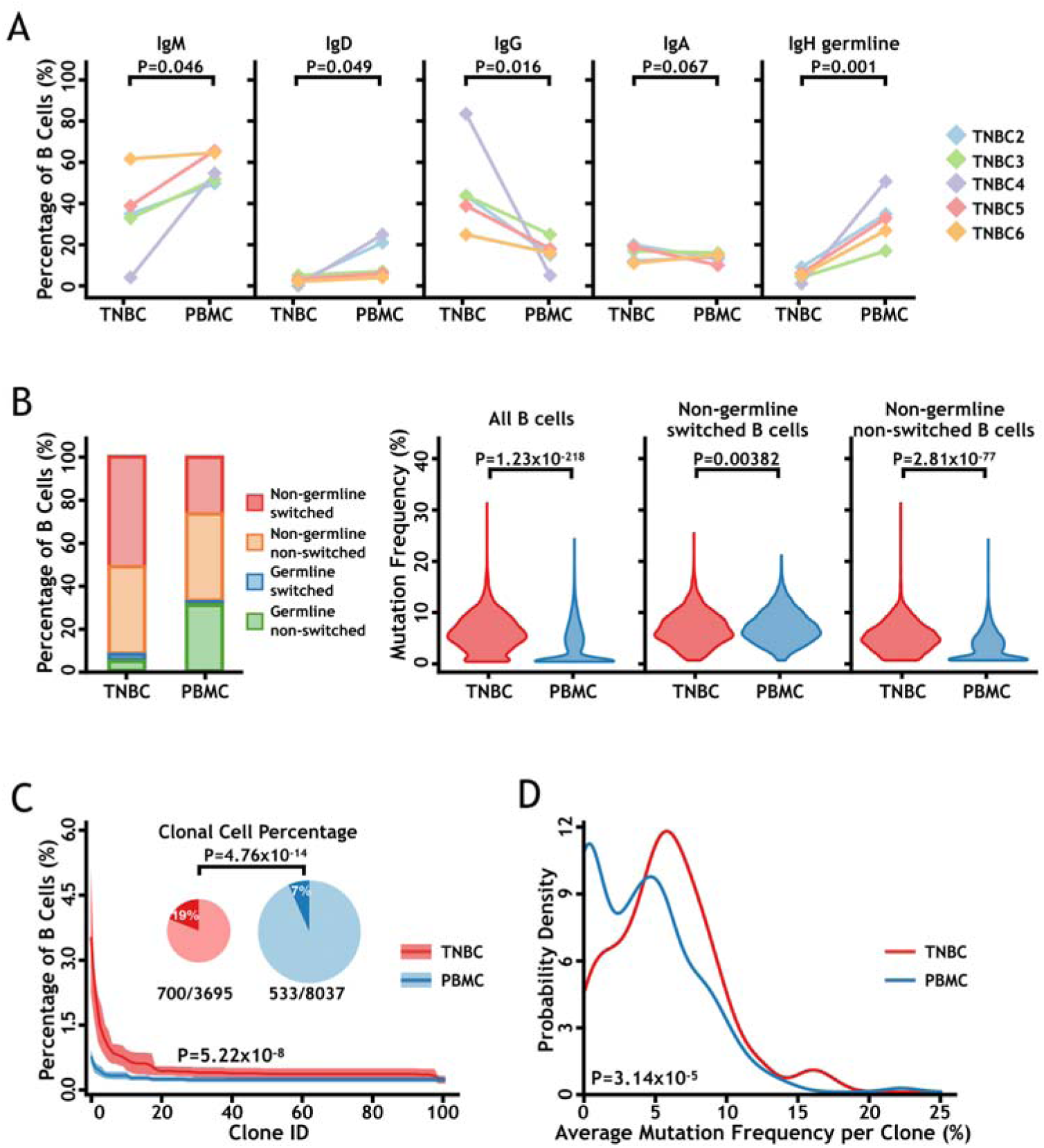
Single cell IgH V(D)J analysis of B cells in tumor and peripheral blood samples from TNBC patients. A. Distributions of IgH isotypes and IgH germline in B cells from TNBC and PBMC samples. The p-values were calculated by student’s t test. B. Comparisons of class switching recombination (CSR) and somatic hypermutation (SHM) between B cells from TNBC and PBMC samples. The left bar-plot presents the percentages of cells that have germline or non-germline and class-switched or non-switched IgH alleles. The violin plots from left to right show the comparisons of SHM rates for all B cells, for non-germline and class-switched B cells, and for non-germline and non-switched B cells. The p-values were calculated by student’s t test. C. TNBC tumors contained more and larger B cell clones than peripheral blood samples. The percentages of clonal B cells in all B cells are presented by upper pie charts. The size distribution of B cell clones showed that TNBC samples have larger B cell clones than PBMC samples. The p-values were calculated by student’s t test. D. TNBC clones have higher IgH mutation frequency than PBMC clones. The average IgH mutation frequencies of each clone are plotted versus the percent of clones with that mutation level. The p-values were calculated by student’s t test.

We could identify individual B cells containing the same rearranged and mutated IgH alleles and also B cell clones within which all B cells share the same germline IgH(Meng et al., 2017). As shown in Fig2C, TNBC infiltrated B cells contained significantly higher percentages of B cell clones (19% vs 7%), which are also larger and more complex than the clones in PBMC (Fig2C and FigS4E). The SHM rates in clones from TNBC are higher than the clones from PBMC (Fig2D). Further identification of the potential tumor antigens for these dominant BCR clones(Sanchez-Trincado et al., 2017) will be the next urgent step to understand the roles of tumor infiltrated B cells in TNBC.

To further dissect the cellular diversity of infiltrated B cells in TNBC, we analyzed 2,526 and 6,106 B cells that have both RNA-seq data and a single assembled productive IgH allele in tumor and peripheral blood samples, respectively. Unsupervised clustering of the combined samples (TNBC2-6 and PBMC2-6) at low resolution revealed 4 clusters with distinct transcriptional signatures (Fig3A-B, FigS5 and Table S4). Cells from different tumor and blood samples contributed to each cluster, suggesting lack of sample batch effect and conserved differentiation process (FigS6). These 4 clusters include naïve B cells (IgM^+^ and IgD^+^), memory B cells (CD27^+^), plasma cells (CD38^+^), and CD14^+^ atypical B cells (CD14^+^). In agree with the BCR results above, B cells infiltrated in TNBC are mostly tissue-resident memory B cells, whereas PBMC samples contain more naïve B cells (Fig3C).

**Figure 3.**
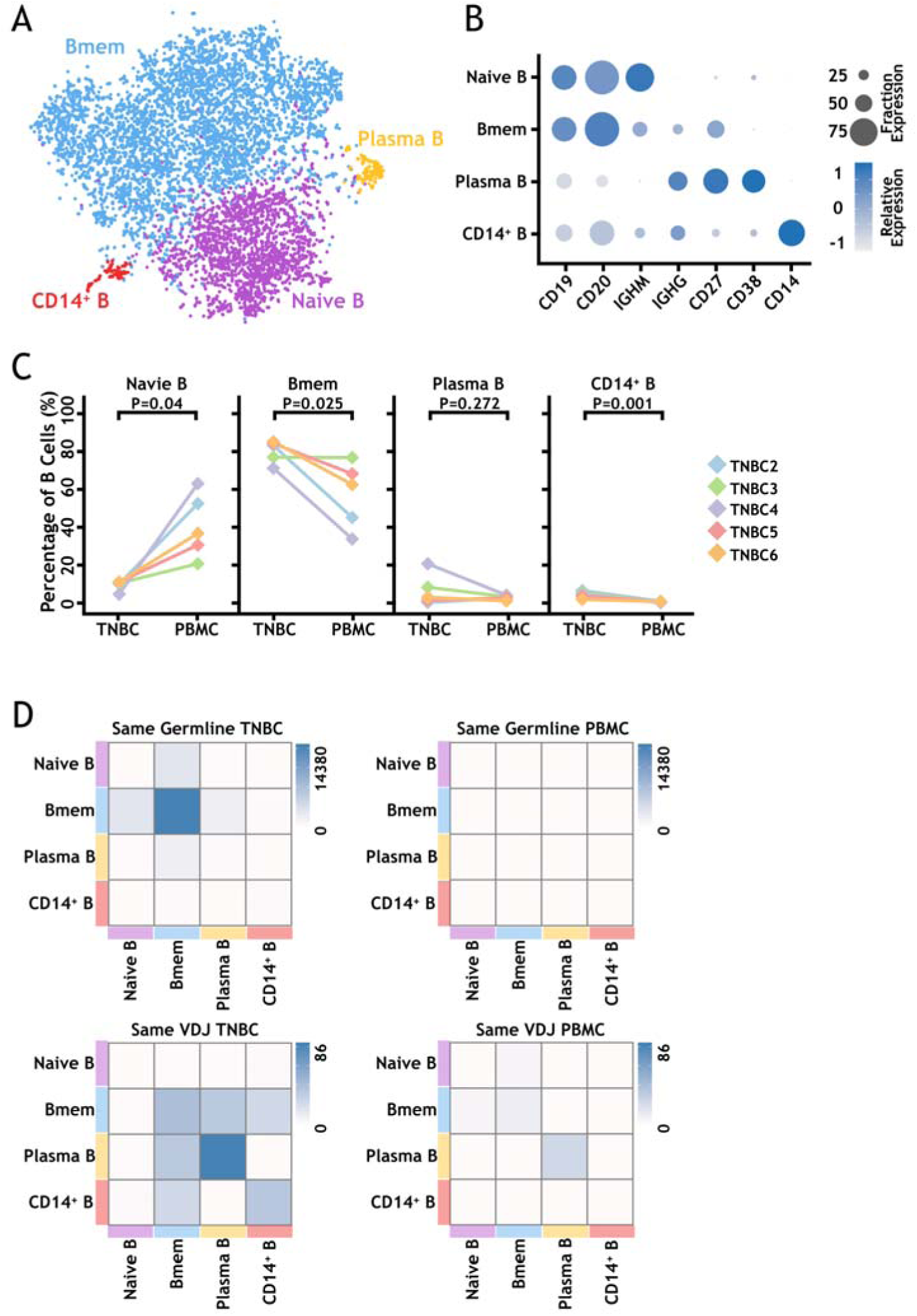
Low resolution of single cell transcriptome analysis and IgH V(D)J lineage analysis of B cells from TNBC patients. A. The t-SNE projection of 8,632 B cells from TNBC patients shows four major cellular clusters. B. Selected marker genes to define each B cell cluster. C. The distributions of four clusters in each patient sample. The p-values were calculated by student’s t test. D. Heat map analysis to show the distribution of cells that share the same IgH germline annotation (upper panel) and the same IgH V(D)J sequences (lower panel) for TNBC and PBMC samples.

With the precise IgH V(D)J and SHM information for each B cell, we were able to trace the proliferation and differentiation of B cells in tumor. After normalization, in a heat map to demonstrate intra- and inter-cluster distribution of B cell clones with the same IgH germline (Fig3D), we found that memory B cells contribute mostly for the IgH BCR clonal trees in TNBC samples, suggesting extensive SHM in those infiltrated memory B cells happening inside tumor. On the other hand, the higher percentages of B cells in TNBC plasma cells contain the same IgH V(D)J.

The higher resolution unsupervised clustering further separated all B cells into 13 clusters with distinct transcriptional signatures (Fig4A-B, FigS5 and Table S5). By combinationally analyzing the expression of known B cell marker genes(Jackson et al., 2008; Kurosaki et al., 2015) and productive IgH sequences, we could annotate these 13 subgroups (Fig4B). They include naïve B cells (C1 and C2), IgM^+^CD27^+^ memory B cells (C3, C4, C6, and C7), IgM^+^CD27^-^ atypical memory B cells (C5), class-switched memory B cells (C8, C9, and C10), plasma cells (C11), germinal center B cells (C12), and CD14^+^ atypical B cells (C13). The distribution of those B cell clusters in tumor samples from different patients showed the larger variation than that in peripheral blood samples (Fig4C).

**Figure 4.**
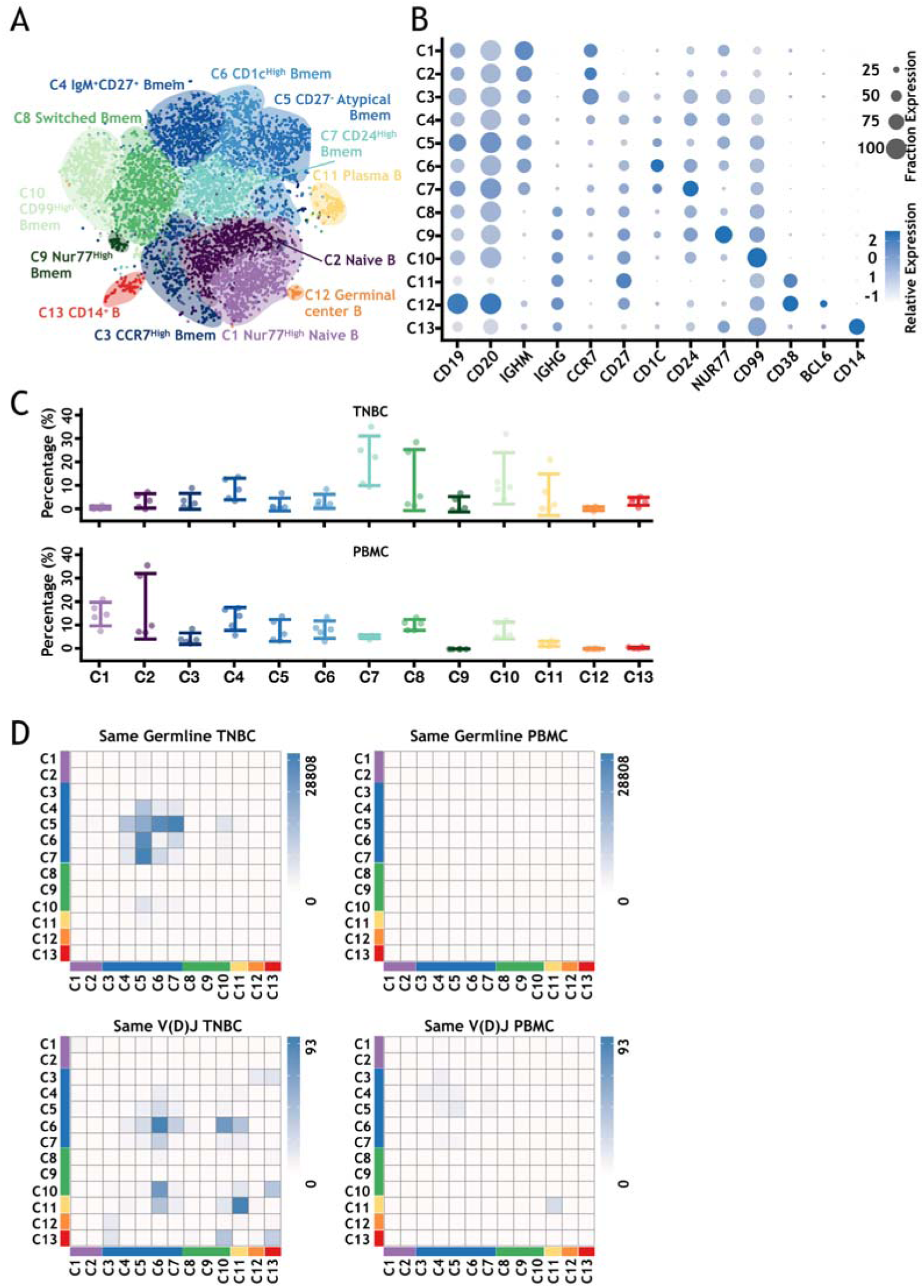
High resolution of single cell transcriptome analysis and IgH V(D)J lineage analysis of B cells from TNBC patients. A. The t-SNE projection of 8,632 B cells from TNBC patients shows thirteen major cellular clusters. B. Selected marker genes to define each B cell cluster. C. The distributions of thirteen clusters in each patient sample (upper panel: TNBC; lower panel: PBMC). D. Heat map analysis to show the distribution of cells that share the same IgH germline annotation (upper panel) and the same IgH V(D)J sequences (lower panel) for TNBC and PBMC samples.

The presence of germinal center B cells (C12), CD1c^High^ memory B cells (marginal zone B cells, C6)(Weller et al., 2004), and various class-switched memory B cell subgroups (C8, C9, and C10) in TNBC tumors suggests the locally ongoing class switching recombination and SHM in potential tertiary lymphoid structures(Colbeck et al., 2017; Dieu-Nosjean et al., 2014) within tumors. In agree with this, we also identified follicular helper T (Tfh) cells in the T cell clusters and various dendritic cell populations in TNBC samples (data not shown). All these results suggested the existence of functionally active germinal centers in TNBC tumor tissues.

Finally, we also tried to identify the potential Breg cells in TNBC. However, although IL10 expression could be well detected in monocytes and macrophages (FigS7), we could not identify specific IL10-expressing B cell populations.

In TNBC, mainly non-switched memory B clusters (C4-C7) were significantly involved in IgH BCR clonal trees (Fig4D). In particular, the CD27^-^ atypical memory B cells (C5) have the highest intra-cluster diversity, suggesting the more ongoing SHM within the cluster. In TNBC, the CD27^-^ atypical memory B cells also shared the same germline significantly with the IgM^+^CD27^+^ B memory cells (C4), the CD1c^High^ B memory cells (C6), and CD24^High^ B memory cells (C7).

In the heat map for the same V(D)J within and between different B cell clusters (Fig4D), plasma cells (C11) have the highest percentage of cells with the same IgH allele within clusters in both TNBC and PBMC, suggesting their clonal proliferation. Interestingly, in TNBC, the CD1c^High^ memory B cells (marginal zone B cells, C6) also show significantly higher clonal expansion than other clusters. In addition, such CD1c^High^ B memory cells share the same IgH allele mostly with the CD99^High^ B memory cells (C10), and also with the plasma cells (C11) and CD24^High^ B memory cells (C7). The CD1c^High^ B memory cells have the most shared the same IgH with other B cell clusters and could be the major direct progenitor cells of these B cells in TNBC.

Recent single-cell analysis of infiltrated T cells in human melanoma revealed a subpopulation of T cells, which is associated with active proliferation and tumor reactivity(Li et al., 2018). In our TNBC samples, infiltrated T cells also showed significant ongoing proliferation evaluated by both cell proliferation signature and cell cycle signature (FigS8). On the contrary, most of TNBC infiltrated B cells, except germinal center B cells (FigS8), are quiescent.

The roles of B lymphocyte infiltration in breast cancer and other types of cancers have been still very controversial (Shen et al., 2018; Wouters and Nelson, 2018). In most of studies, the presence of infiltrated B lymphocytes was determined by IHC of pan-B cell markers such as CD20. We hypothesized that distinct infiltrated B cell clusters might have diverse functions in tumor immunology and the combinational transcription signature based on all their differentially expressed marker genes (Table S6) for individual B cell clusters might be a better indication of their presence in tumors. In deed, using expression data and clinical information of TNBC patients from the METABRIC consortium, we found that the transcription signatures of the naïve B cells, memory B cells, and C2 naïve B cells are significantly associated with improved overall and disease-free survival of TNBC patients (Fig5A). More importantly, comparing with classic single B cell markers such as CD19 and CD20, those B cell signatures show much stronger hazard ratios (HRs) in both univariable and multivariable analyses for TNBC patients (Fig5B and FigS9), suggesting that they provide better prognostication that CD19 or CD20 alone. Using available data from TCGA consortium, the memory B cell signature also demonstrated more significant association with overall survival and stronger HRs for patients with cervical squamous cell carcinoma and endocervical adenocarcinoma (CESC), sarcoma(SARC), skin cutaneous melanoma (SKCM) and uterine corpus endometrial carcinoma (UCEC) (FigS10 and Table S7).

**Figure 5.**
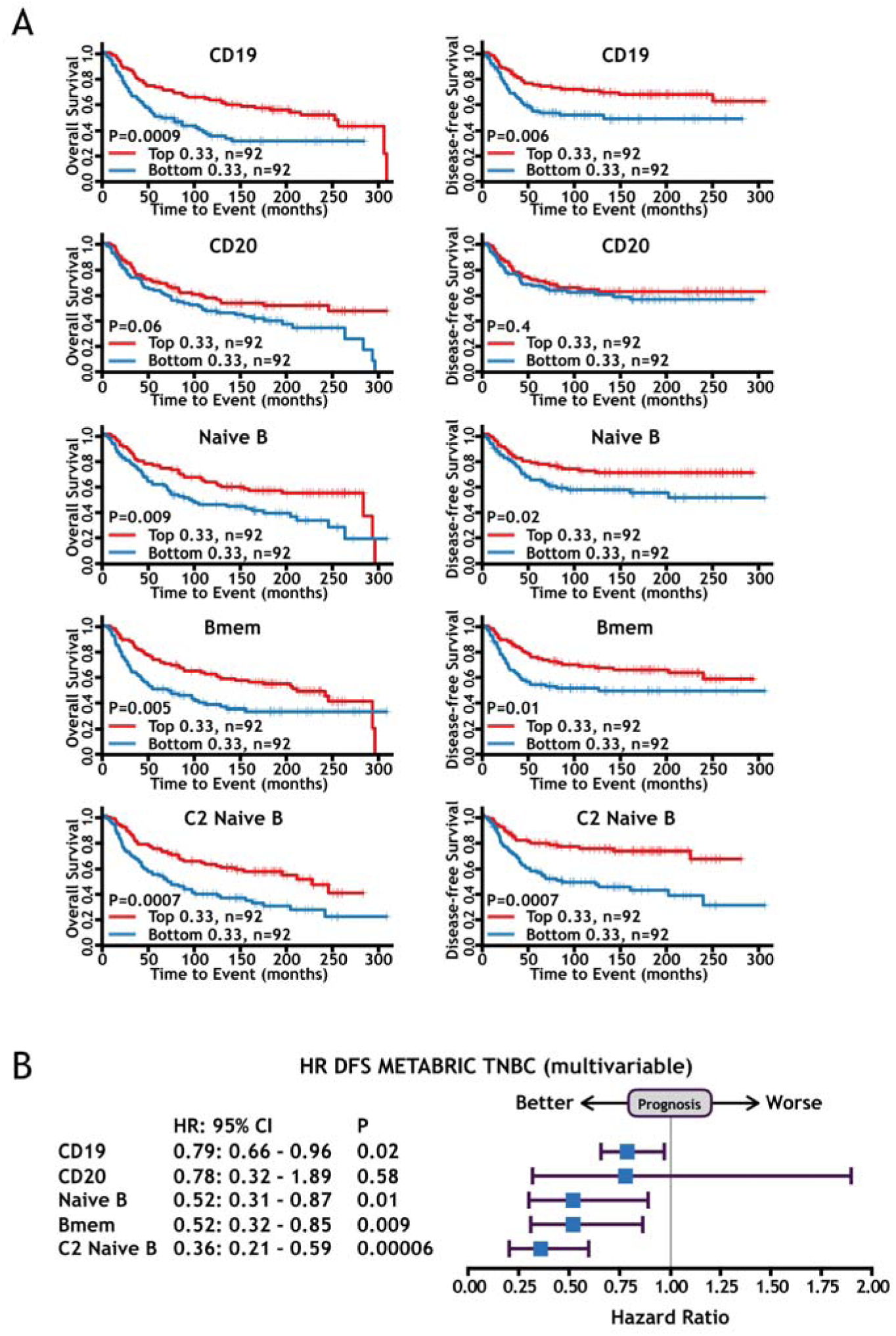
The B cell signatures based on single-cell RNA-seq results are significantly associated with improved survival for TNBC patients and provide better prognostication than classic single B cell markers. A. Kaplan–Meier survival curves for overall survival (left panel) and disease-free survival (right panel) of TNBC patients from the METABRIC consortium according to single gene expression (CD19 and CD20) and combined gene transcription signatures (Naïve B cells, memory B cells, and C2 Naïve B cells). The p-values were calculated by log-rank test. B. Prognostic effect of CD19, CD20, Naïve B, memory B and C2 Naïve B signatures in TNBC patients from the METABRIC consortium. Forest plots show HRs (blue squares) and confidence intervals (horizontal ranges) derived from Cox regression survival analyses for disease-free survival in multivariable analyses adjusted for lymph nodes status, tumor size, age at diagnosis, and histological grade.

It has been widely accepted that the immune microenvironment plays essential roles in tumor development and therapy(Finn, 2018; Wei et al., 2018). While T cells have been mostly studied and therapeutically targeted in tumor immunology, the roles of B cells have been only noticed recently(Liu et al., 2018; Sarvaria et al., 2017; Shen et al., 2018; Tsou et al., 2016; Wouters and Nelson, 2018). However, the results on association of tumor infiltrated B cells with the prediction and prognosis of various tumors, such as breast cancer, have been very controversial(Shen et al., 2018), which might be due to the heterogeneity of both breast cancer patients and B cell subtypes. We focused on TNBC, the most immunogenic breast cancer subtype, and dissected the tumor infiltrated B cell populations by paired single-cell antigen receptor repertoire and whole-transcriptome sequencing. We presented a comprehensive single-cell analysis of TNBC infiltrated B cells and found that most of TNBC infiltrated B cells showed more mature and memory B cell characteristics. These TNBC infiltrated memory B cells have higher clonality and extensive IgH class switching recombination and somatic hypermutations, likely happened within tumor and associated with tumor antigen experience. Our results also suggested the existence of functionally active germinal centers in TNBC tumor tissues. On the other hand, we could not detect obvious Breg populations characterized by IL-10 expression in TNBC. The B cell signatures based on single-cell RNA-seq results significantly associated with improved survival for TNBC patients and provide better prognostication than classic B cell markers (CD19 or CD20).

Precisely targeting the immune subtype cells will lead to better therapeutic efficacy and also fewer side effects at the same time. Dissecting the heterogeneity of tumor infiltrated B cell subgroups provides the essential information to functionally reveal their roles in development and treatment of TNBC. B cells could positively participate in tumor immunology through antibody production, antigen-presentation, cytokine and chemokine production, and other immunoregulation mechanisms(Liu et al., 2018; Sarvaria et al., 2017; Tsou et al., 2016). On the other hand, B regulatory cells (Bregs) play important roles in maintaining immune homeostasis mainly by secreting IL-10 or directly interact with target cells(Rosser and Mauri, 2015). Dissection of the mechanisms regulating the origin and dynamic distribution of tumor infiltrated cell populations will provide new clues for tumor immunotherapy. Further studies are needed for the mechanisms leading to the clear association of infiltrated B memory cells with survival of TNBC patients. Pathways that regulate the infiltration, expansion, and differentiation of memory B cells could be used as new targets for TNBC immunotherapy.

Although we only thoroughly analyzed TNBC infiltrated B cells here, we also obtained single cell transcriptional data for more than 88,254 other immune cells and single cell TCR results for more than 58,095 T cells (GSE123192). These data will be invaluable to understand the overall tumor immune environment and potential interactions between different immune cell types, as well as to identify new immunotherapy targets. Further studies on the characterization of all infiltrated immune cell types in TNBC and mechanisms how TNBC escaped from immunosurveillance in human patients will provide essential guidance for immunotherapy of TNBC. Our analysis and experimental strategies could also be widely applied to B cell analysis in other tumor types.

## Methods

### Reagents and antibodies

Antibodies used in this study included PE anti-human CD45 (BioLegend, Cat#304008), PerCP-Cy^TM^5.5 Mouse Anti-Human CD56 (BD, Cat#560842), Anti Human CD11b PE-Cyanine7 (eBioscience, REF 25-0118-42), APC anti-human CD3 (BioLegend, Cat#300312), FITC anti-human CD20 (BioLegend, Cat#302304) and FITC anti-human CD19 (BioLegend, Cat#302206).

### Patient samples

This study was approved by the human research ethics committee of the Xingxiang central hospital (Xinxiang, P.R. China). All participated patients provided written informed consent. The breast cancer tissues were collected during surgery from patients who had not experienced any chemotherapy and other treatments. Peripheral bloods were collected from the same patients.

### Isolation of lymphocytes from peripheral blood and breast cancer biopsies

Peripheral blood mononuclear cells (PBMCs) were isolated using Lymphoprep^TM^ (Sigma-Aldrich) solution according to the manufacturer’s instructions. Briefly, 3 mL of fresh peripheral blood was 1:1 diluted with 0.9%NaCl and carefully loaded over 3ml Lymphoprep in a 15 ml falcon tube.

After centrifugation at 800g for 30 minutes at room temperature, the mononuclear cells (at the sample/lymphoprep interface) were carefully transferred to a new tube and washed with 1x PBS at 250g for 10 minutes. These lymphocytes were resuspended with FACS sorting buffer (PBS supplemented with 2% fetal bovine serum (FBS, Sciencell)) and stained with indicated FACS antibodies. The hCD45 positive cells were sorted using BD FACSAria II.

Fresh breast tumor samples were cut into small pieces and gently triturated with a 5 mL syringe plunger on a 70 μm Cell-Strainer (BD) in RPMI-1640 medium (Invitrogen) with 2% FBS on ice until uniform cell suspensions were obtained. The cells were subsequently passed through cell strainers and centrifuged at 400g for 10 min. The cell pellets were resuspended in 6 ml RPMI-1640 medium supplemented with 2% FBS. The next steps were the same as peripheral blood sample preparation.

### Whole-exome sequencing for germline and somatic mutations in breast cancer patients

Genomic DNAs isolated from peripheral blood and tumor tissues were subjected to whole-exome library construction and 150bp paired-end sequencing. The sequencing reads were aligned to the human reference genome (hg19) using BWA(Li and Durbin, 2010) and germline and somatic SNV/indels were called using Varscan2(Koboldt et al., 2012). Germline SNV/indels were called by “java -jar VarScan.v2.3.7.jar mpileup2cns--min-avg-qual 20 --output-vcf --variants 1” and then annotated by ANNOVAR(Wang et al., 2010). Variations recorded as pathogenic in ClinVar were listed in Table S3. The stop-gain/loss, indel, and nonsynonymous mutations with less than 1% in ExAC_ALL population and ExAC_EAS population were kept; and for nonsynonymous mutations, sift score <0.02 and polyphen score >0.9 were required (Table S3). Variations in cancer related genes were selected manually from Table S3 and shown in FigS2A. Somatic SNV/indels were called by “java -jar VarScan.v2.3.7.jar somatic --output-vcf 1--min-coverage 20 --strand-filter 1 --min-var-freq 0.1”, and somatic SNV/indels with at least 3 variant reads in tumor and less than 5% of variant reads in blood were kept (Table S3) for further annotation by ANNOVAR(Wang et al., 2010) and signature analysis by MutationalPattern(Blokzijl et al., 2018). Somatic variations with less than 1% in ExAC_ALL population and ExAC_EAS population, sift score <0.02 and polyphen score >0.9 were selected to generate the heatmap in FigS2B.

### Single-cell RNA sequencing and TCR/BCR library construction

The FACS-sorted viable hCD45+ cells were counted with Trypan blue using hemocytometer. Cells were then resuspended at 5×10^5^–1×10^6^ cells/ml with a final viability > 85%. Single-cell library preparation was carried out using 3’ library v2 or 5’ V(D)J and gene expression platform as per 10 × Genomics protocol (10 × Genomics, Pleasanton, CA, USA). Cell suspensions were loaded onto a Chromium Single-Cell Chip along with the reverse transcription master mix and single cell gel beads, aiming for 2,000–10,000 cells per channel. Following generation of single-cell gel bead-in-emulsions (GEMs), reverse transcription and amplification were performed. Then amplified cDNAs were purified and sheared. Sequencing libraries were generated with unique sample index for each sample. Libraries were sequenced by Illumina HiSeq X10 or NovaSeq.

### Quality control and filtering of single-cell RNA sequencing data

The 10x Cell Ranger package (version 2.1.1, 10x Genomics) was used to de-multiplex cellular barcodes, to map reads to the hg38 reference assembly (v1.2.0, 10x Genomics), and to generate gene UMI (unique molecular identifier) counts versus cell barcode matrix. The 20 single-cell RNA-sequencing libraries were sequenced to average 74,161 (40,341-191,921) paired-end reads per cell, and average 88.3% (80.2-94.7%) sequencing saturation (Table S1). To remove multiple captures, which is a major concern in microdroplet-based experiments, we excluded top N cells with the highest pANN score calculated by DoubletFinders (McGinnis et al., 2019) for each library separately, where N is inferred from the doublet rates in 10x platform(Zheng et al., 2017b). From the supplementary Fig1a(Zheng et al., 2017b), we can infer the doublet cell number as (0.000879*N+0.702)*0.01*N, where N represents the cell number detected by cell ranger. Cells with the numbers of detected genes < 200 or >5,000 were also deleted. Cells with >50% of the UMI counts belonging to mitochondrial genes were also deleted. Finally, 112,938 cells from total 20 libraries remained for downstream analysis, with 4,811 median UMI per cell (2,442-7,391 for each library, 2,605 and 3,354 for the two 3’ libraries) and 1,600 median genes per cell (1,012-2,267 for each library, 1,012 and 1,151 for the two 3’ libraries).

### Canonical Correlation Analysis, Dimensionality Reduction and Clustering

For all cells after above quality control steps, the UMI cell barcode matrix of each library were processed by Seurat (version2.3.4) (Butler et al., 2018) for normalization, dimension reduction, batch effect removement, graph-based clustering, cluster specific marker genes detection, and visualizations. For each library, library-size normalization to each cell was done by NormalizeData; the variability of the numbers of UMIs was regressed out by ScaleData; the variable genes were calculated by FindVariableGenes. Then all 20 libraries were combined together using RunMultiCCA. The genes used for correlation components (CCs) calculation were the combination of top 2000 dispersion genes for each library (total 3,175 genes, at least appeared in two samples). The cells were then aligned with AlignSubspace using 10 CC dimensions. The FindClusters was used to cluster cells using the 10 aligned CCs at resolution 0.1 (total 4 clusters). The clustering results were visualized with t-distributed stochastic neighbor embedding (tSNE) dimensionality reduction using RunTSNE (10 aligned CCs) and TSNEPlot.

### B cell clustering analysis

B cells from TNBC2-6 and PBMC2-6 were selected for sub-grouping when the cell barcode had a single productive IgH BCR expressing in the BCR library and had no TCR expressing in the TCR library. Then the UMI cell barcode matrices of B cells were subjected to NormalizeData, ScaleData, and FindVariableGenes as above. RunMultiCCA was used to combine the B cell from all libraries, and 10 CC dimensions were used for AlignSubspace. The FindClusters was used to cluster cells using the 10 aligned CCs at resolution 0.1 for low resolution grouping (Fig3) and 1.0 for high resolution grouping (Fig4). The clustering results were visualized with t-distributed stochastic neighbor embedding (tSNE) dimensionality reduction using RunTSNE (10 aligned CCs) and TSNEPlot. Marker genes for each cluster were detected by FindAllMarkers function, except setting min.pct=0.25. Genes were ranked by their fold-change (from largest to smallest) and the top10 genes of each cluster were showed in the heatmap (FigS5).

We defined signature genes of each cell group for survival analysis as described previously(Savas et al., 2018). Cells from TNBC tissue were used for signature gene define by DECENT(Ye et al., 2019) using default parameters without spike-ins, and cutoff for signature genes was fold-change> 2 and FDR (Benjamini–Hochberg adjusted P value) < 0.01. As cells in each group are from 5 different samples, we attempted to correct for possible batch effects by including a dummy batch variable in the model for the combined data when using DECENT.

### B Cell Receptor (BCR) Analysis

The 10x Cell Ranger package (version 2.1.1, 10x Genomics) was used to assemble raw sequencing reads of each BCR library into contigs that represent the best estimate of transcript sequences in each cell barcode. Then the assembled contigs (filtered_contig.fasta files from Cell Ranger) were processed by IMGT/HighV-QUEST(Alamyar et al., 2012) to assign V(D)J germline segments to each contig using default settings, except setting “species” to “Homo sapiens” and setting “Receptor type or locus” to “IGH”. The output files of IMGT/HighV-QUEST were analyzed by Change-O(Gupta et al., 2015) and custom scripts to select cells with a single productive IgH. Productive IgH was selected using ParseDb.py with parameters -f FUNCTIONAL -u T. If there were more than one productive IgH detected within one cell, the most abundant IgH sequence was kept when the UMI count of most abundant IgH sequence is > 10-fold of the second most abundant IgH sequence. We obtained 11,732 cells with single productive IgH and 8,632 cells with both transcriptome and IgH sequence for TNBC2-6 and PBMC2-6 samples (Table S2). IgH sequences were considered clonal when V, D, J usage and the CDR3 sequence length were the same within groups of B cells. SHM number and frequency were calculated using V and J regions (D region and junctions were not included). Lineage trees were inferred using PHYLIP version 3.69(Felsenstein, 1989).

### Survival analyses

RNA-Seq gene expression profiles (Level 3) and clinical data for all tumor types were downloaded from the TCGA data portal (https://gdc-portal.nci.nih.gov/). The METABRIC gene expression profiles and clinical data were downloaded from the cBioPortal website (http://www.cbioportal.org/study?id=brca_metabric#summary). The TNBC samples were defined by IHC results with HER2^-^, ER^-^, PR^-^, and samples with normal subtype by PAM50 were removed. Samples were sorted by expression level of gene or gene set, and top 33% versus bottom 33% of samples were used to generate survival curves. Multivariable analyses were adjusted by lymph nodes status, tumor size, age at diagnosis, and histological grade. For survival analysis and Hazard Ratio calculation, R package survival(Therneau and Grambsch, 2013) was used.

### Paired BCR and single-cell RNA sequencing data analysis

To trace the proliferation and differentiation of different B cell clusters, we counted the same germline event number and same V(D)J event number within and between each B cell clusters. The same germline event was counted when two cells were in the same clonal B cell tree, and the same V(D)J event was counted when two cells have the same observed IgH sequences. The same germline event and the same V(D)J event could be grouped into PBMC (when two cells were both from PBMC samples), TNBC (when two cells were both from TNBC samples), or across PBMC/TNBC (when one cell was from TNBC samples and the other cell was from PBMC samples). Then heatmap of same germline or same V(D)J can be generated for PBMC, TNBC, or PBMC/TNBC using the above event number, which was then normalized by dividing the coordinate cell numbers in each B cell cluster in PBMC samples (for PBMC heatmap), in TNBC samples (for TNBC heatmap) and in all samples (for PBMC/TNBC heatmap).

### Cell cycle analysis

Cell cycle scores were calculated as the percentage of cell cycle gene UMIs out of total UMIs in a cell(Li et al., 2018). We also used CellCycleScoring function of Seurat package(Butler et al., 2018) to calculate the S phase score and G2/M score. The cell was considered to be in S phase when the S phase score >0.1 and > G2/M score. The cell was considered to be in G2/M phase when the G2/M phase score>0.1 and > S phase score.

## Data availability

Raw fastq files for single-cell sequencing and whole exon sequencing have been deposited at the Gene Expression Omnibus (GEO) and Sequence Read Archive (SRA) under accession numbers GSE123192.

## Supplemental information

Supplemental information includes 10 figures and 7 tables.

**Figure S1.**
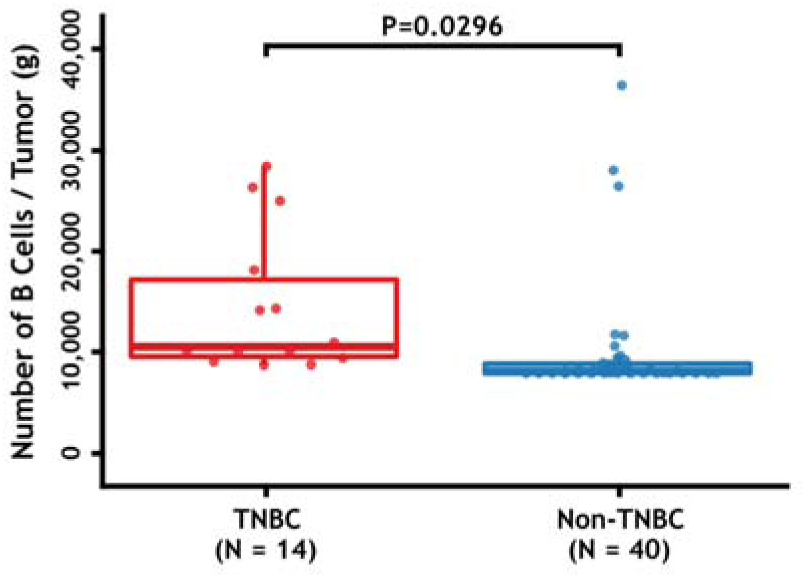
TNBC tumors have more infiltrated B cells than other breast cancer types. The numbers of B cells in tumor samples quantified by FACS were normalized by tumor weight (per gram). The p-values were calculated by student’s t test.

**Figure S2.**
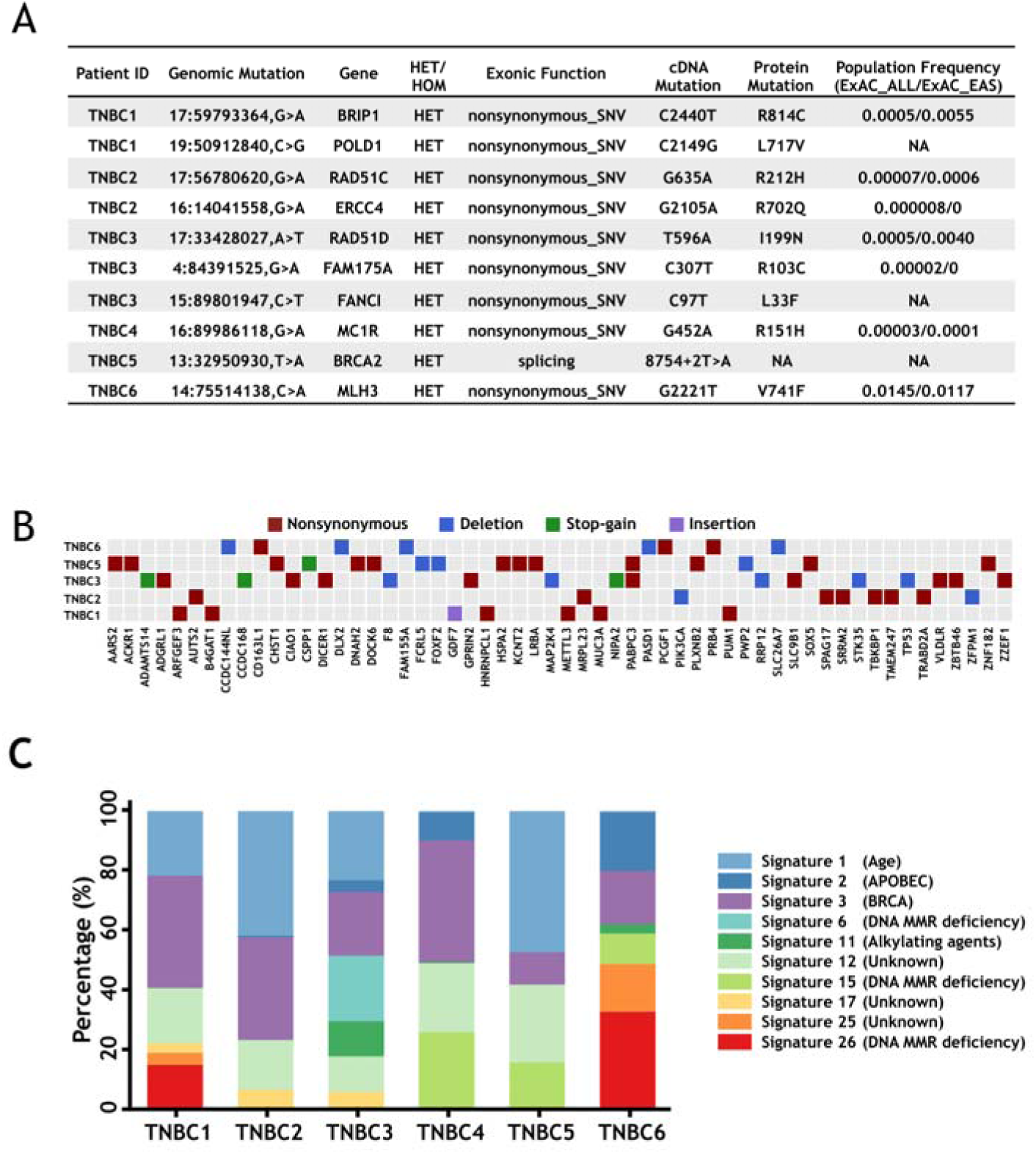
Germline and somatic mutations identified in TNBC patients. A. Potential tumor driver germline mutations were identified in TNBC patients by whole-exome sequencing. B. Somatic mutations in cancer related genes were also identified in TNBC tumor samples. C. The signatures of somatic mutations in TNBC tumor samples.

**Figure S3.**
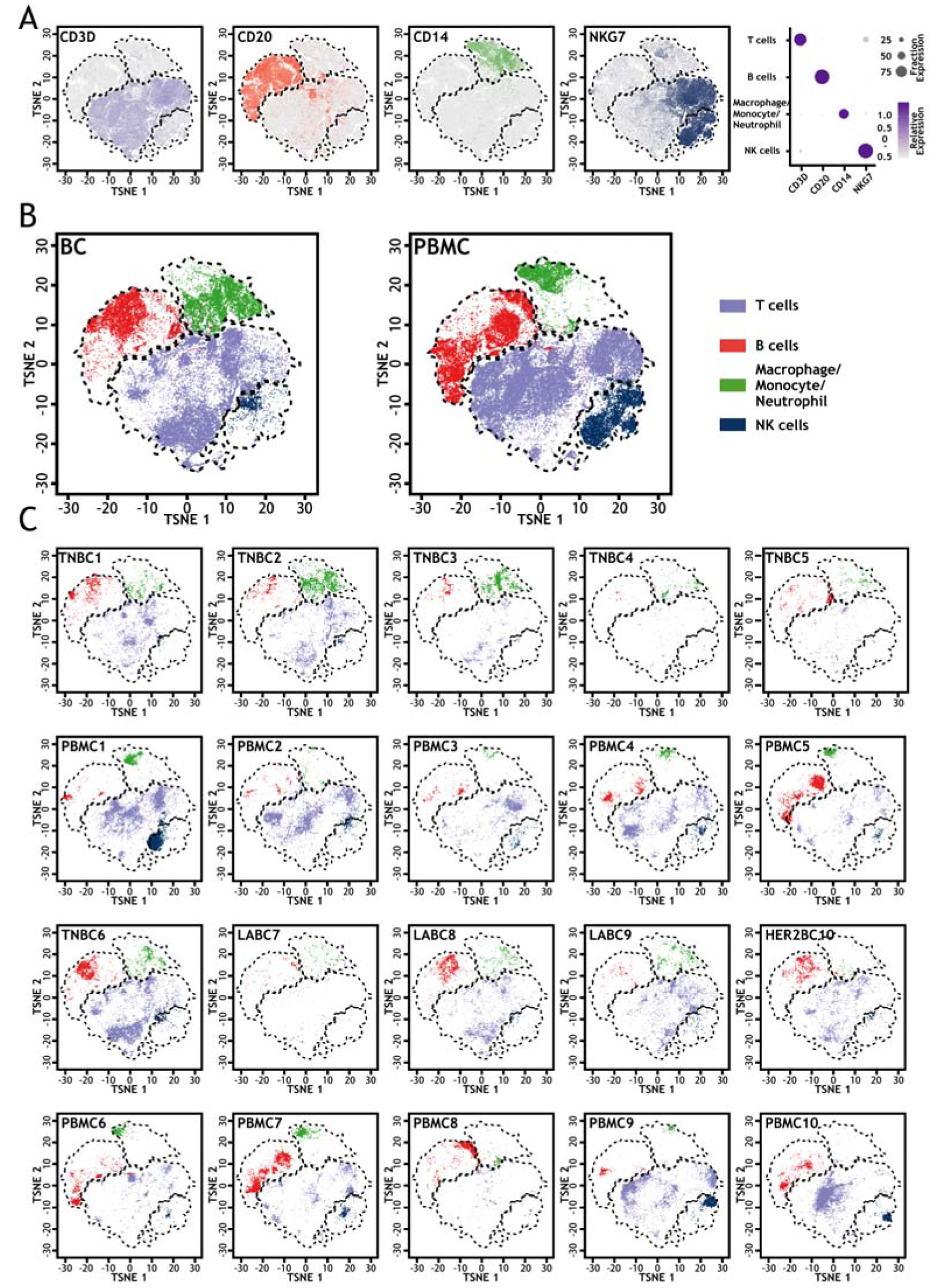
Unsupervised clustering at low resolution on all sequenced samples revealed four major cellular clusters including T cells, B cells, NK cells, and macrophage/monocyte/neutrophil. A. The t-SNE projections of marker genes for T cells (CD3D), B cells (CD20), NK cells (NKG7), and macrophage/monocyte/neutrophil (CD14). B. Both BC and PBMC samples contributed similarly to each cluster. C. Cells from different samples contributed similarly to each cluster.

**Figure S4.**
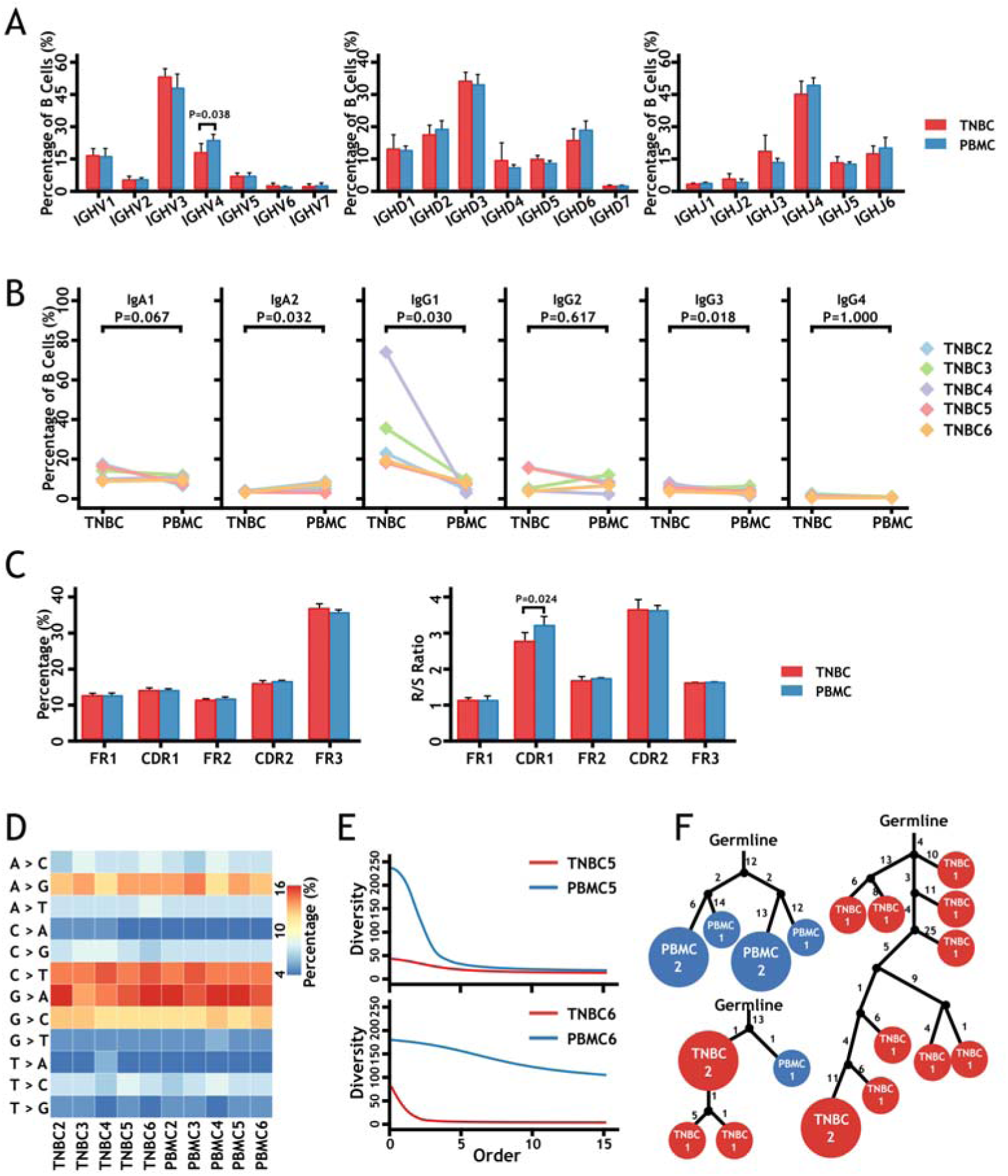
Single cell IgH V(D)J analysis of B cells in tumor and peripheral blood samples from TNBC patients. A. VH, DH, and JH usages in TNBC (red) and PBMC (blue) B cells. B. Distributions of IgH isotypes in TNBC and PBMC samples. The p-values were calculated by student’s t test. C. Comparisons of SHM in different IgH V(D)J regions (Left panel, mutation distributions in different V(D)J regions; Right panel, R/S (Replacement mutations / Silent mutations) ratio comparisons of different V(D)J regions. D. IgH SHM signatures of each sample. E. The IgH diversities of TNBC5 and TNBC6 samples were lower than corresponding PBMC samples. Diversity of clones with at least 2 cells was plotted at different orders (Hill numbers). At an order of 0, the diversity is the number of different clones(Meng et al., 2017). F. Examples for B cell clonal trees with only PBMC cells, only TNBC cells, and both PBMC and TNBC cells. The trees are rooted in the closest IgH V(D)J germline allele in the IMGT database. Numbers indicate somatic mutations. Red circles represent TNBC infiltrated B cells and blue circles represent PBMC B cells. The sizes of the circles are proportional to cell numbers (indicated in the circles, too). Black dots indicate inferred nodes.

**Figure S5.**
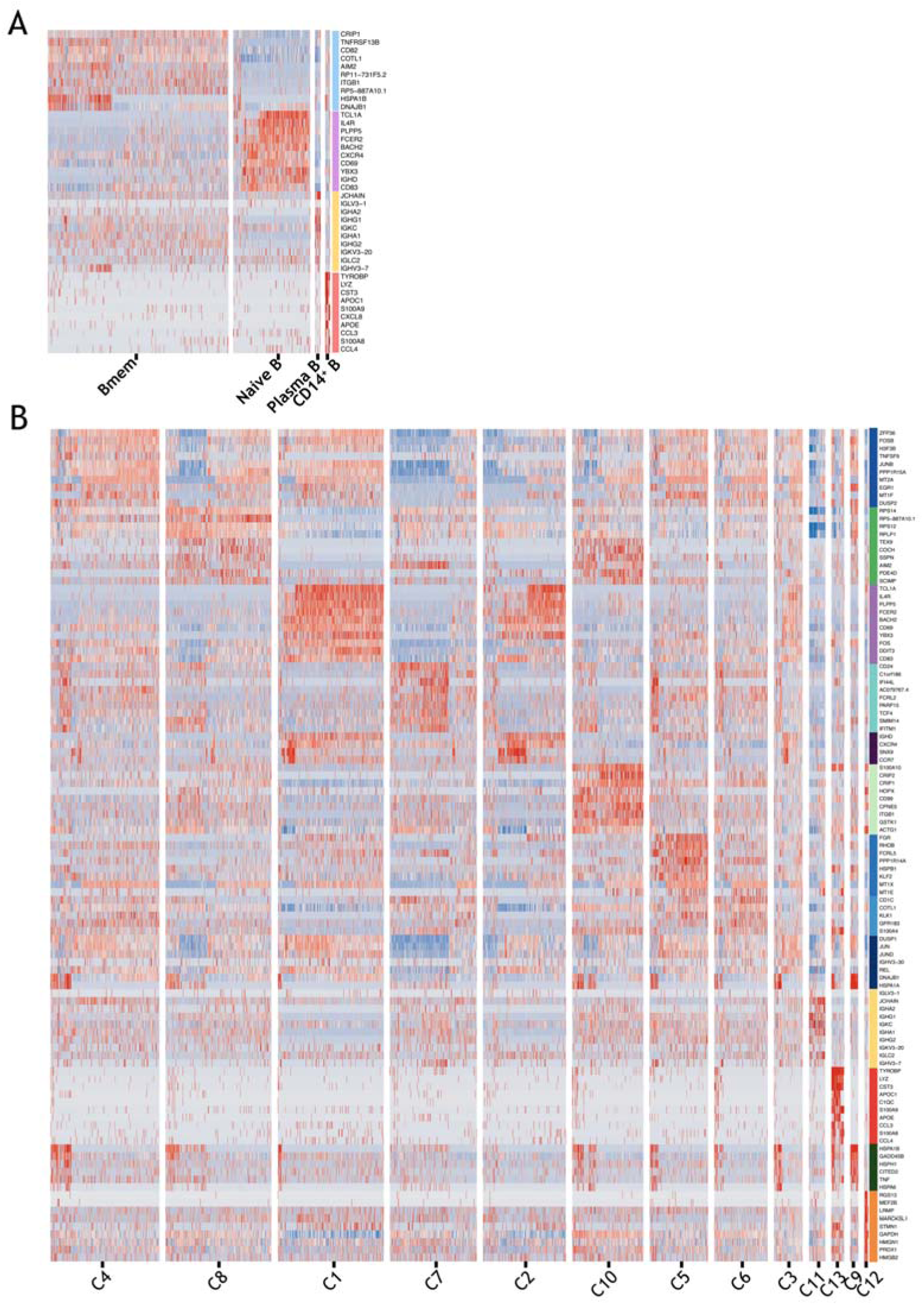
Heat map of marker genes in each B cell group. A. Heat map of top10 marker genes for the four B cell clusters from low-resolution grouping. Gene names are labeled alongside. B. Heat map of top10 marker genes for the thirteen B cell clusters from high-resolution grouping. Gene names are labeled alongside.

**Figure S6.**
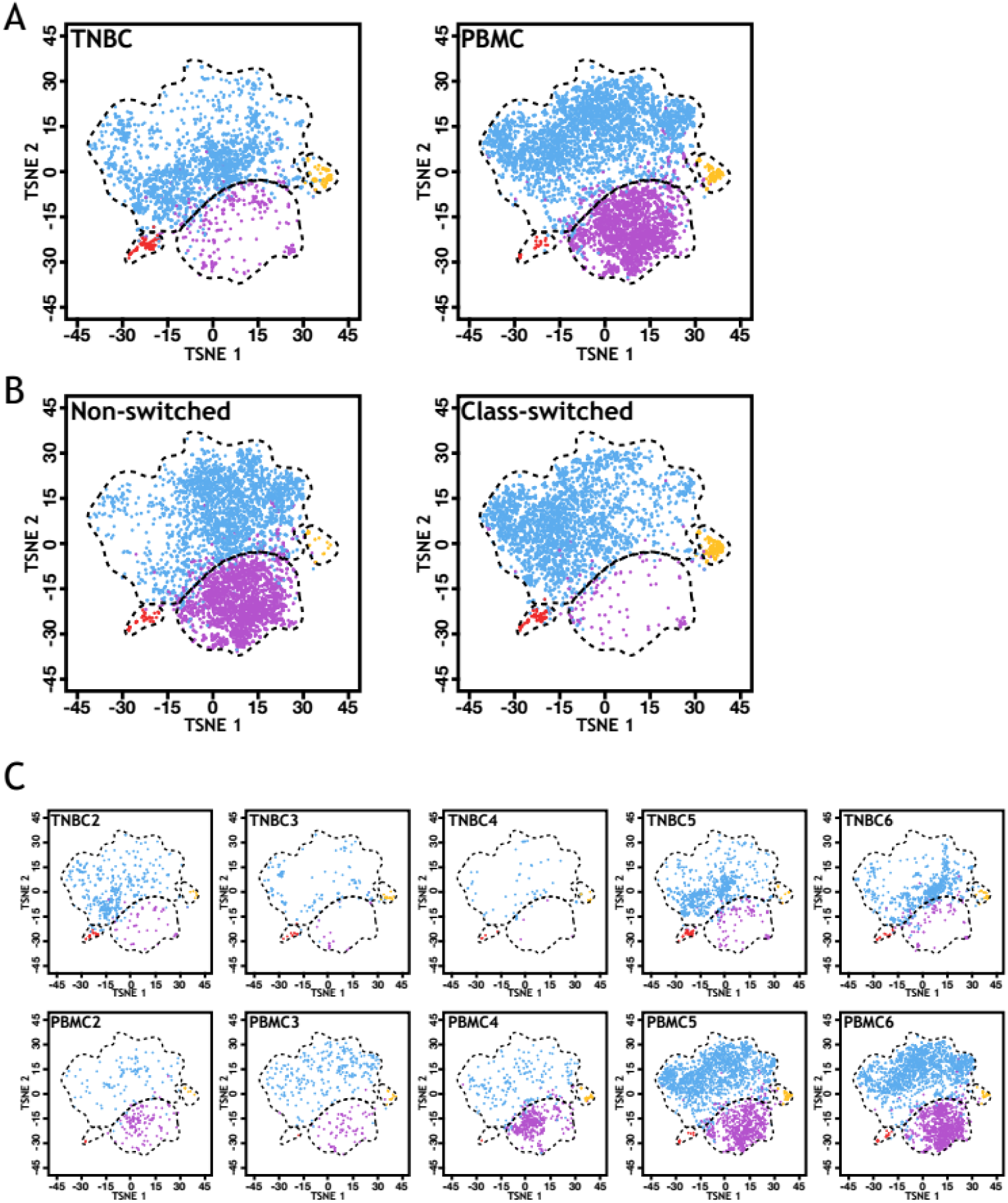
Unsupervised clustering of B cells revealed four different cell clusters. A. Both TNBC and PBMC samples contributed similarly to each cluster. B. IgH class-switched and non-switched B cells could be well separated in t-SNE plots. C. B cells from different samples contributed similarly to each cluster.

**Figure S7.**
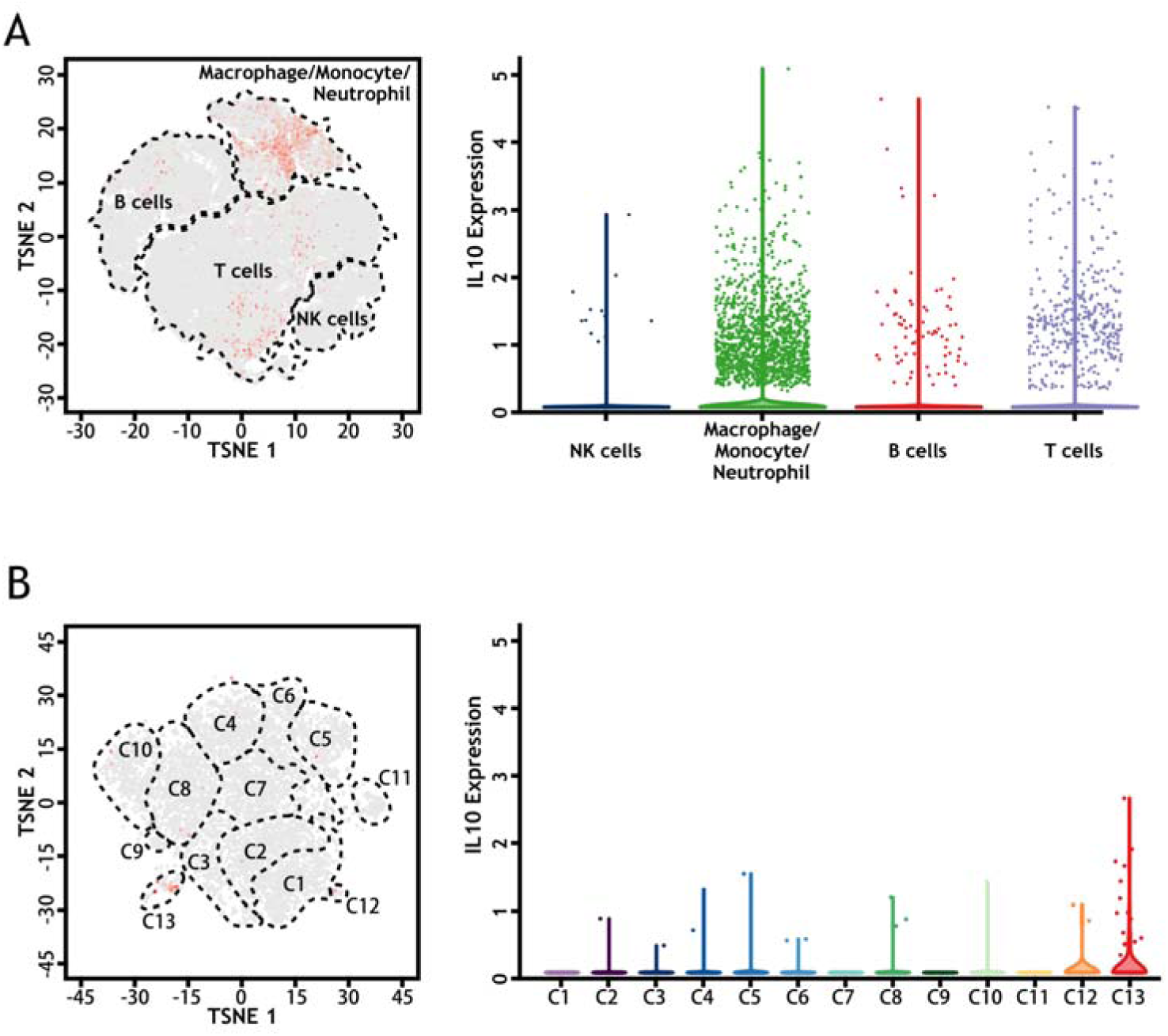
Lack of IL10-expressing Breg cells in TNBC. A. The t-SNE projection (left panel) and violin plot (right panel) of IL-10 expression in CD45+ cells from all patient samples. In the t-SNE plot, each dot represents a cell and the red color represents cell expressing IL-10. In the violin plot, the color represents cell types, the same as in Fig1B. B. The t-SNE projection (left panel) and violin plot (right panel) of IL-10 expression in B cells from TNBC patient samples. In the t-SNE plot, each dot represents a cell and the red color represents cell expressing IL-10. In the violin plot, the color represents cell types, the same as in Fig4A.

**Figure S8.**
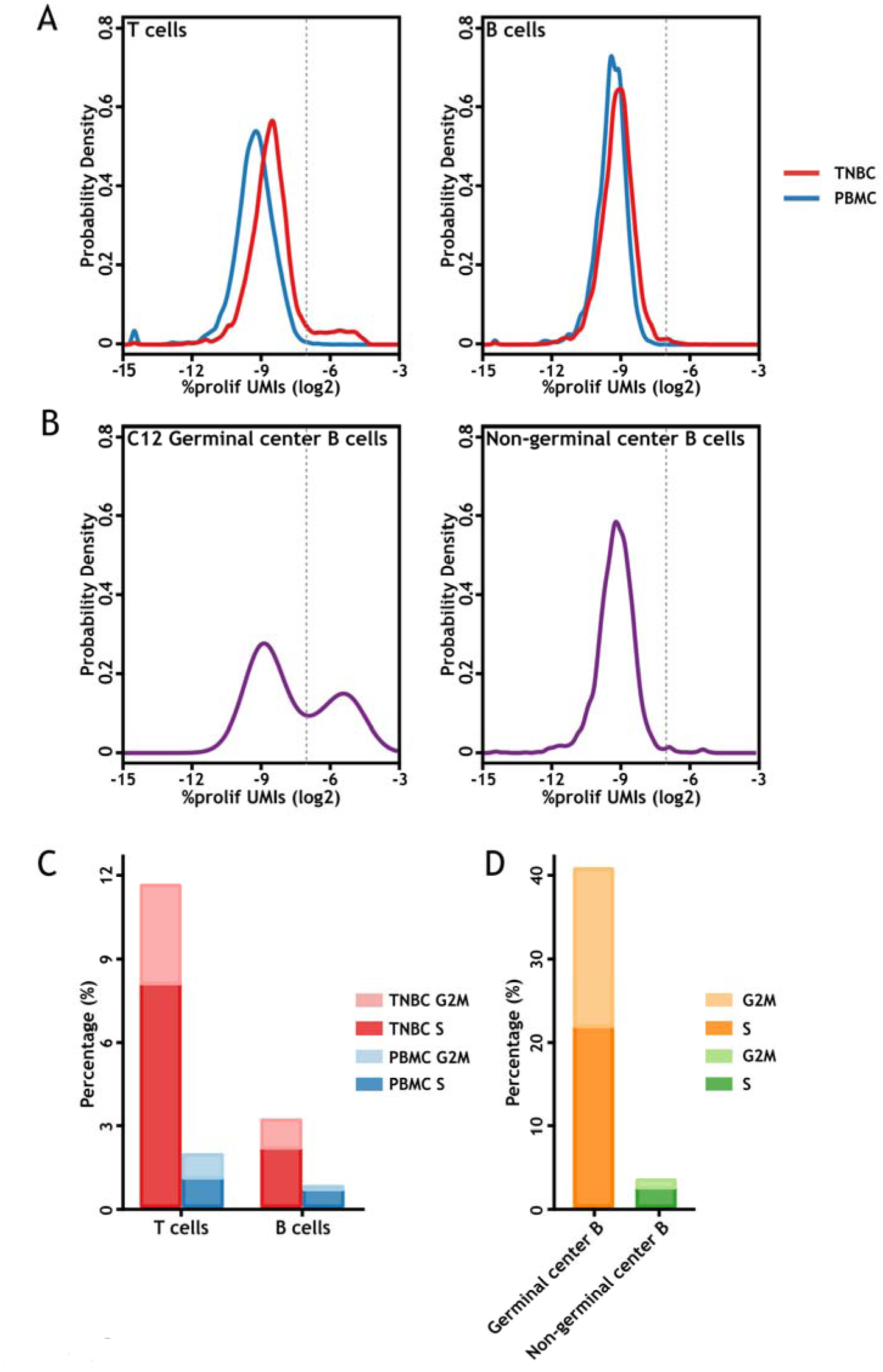
Cell cycle analysis of infiltrated B cells and T cells in TNBC samples. A. Frequency distribution of cell cycle scores on T cells (left panel) and B cells (right panel) in TNBC tumors and PBMC samples. Cell cycle scores were calculated as the percentage of cell cycle gene UMIs out of total UMIs. B. Frequency distribution of cell cycle scores on cluster 12 germinal center B cells (left panel) and non-germinal center B cells in TNBC samples. C. Percentage of proliferating cells (S and G2M) for T cells and B cells in TNBC samples and PBMC samples. D. Percentage of proliferating cells (S and G2M) for C12 Germinal center B cells and Non-germinal center B cells. Data are from 5 TNBC patients.

**Figure S9.**
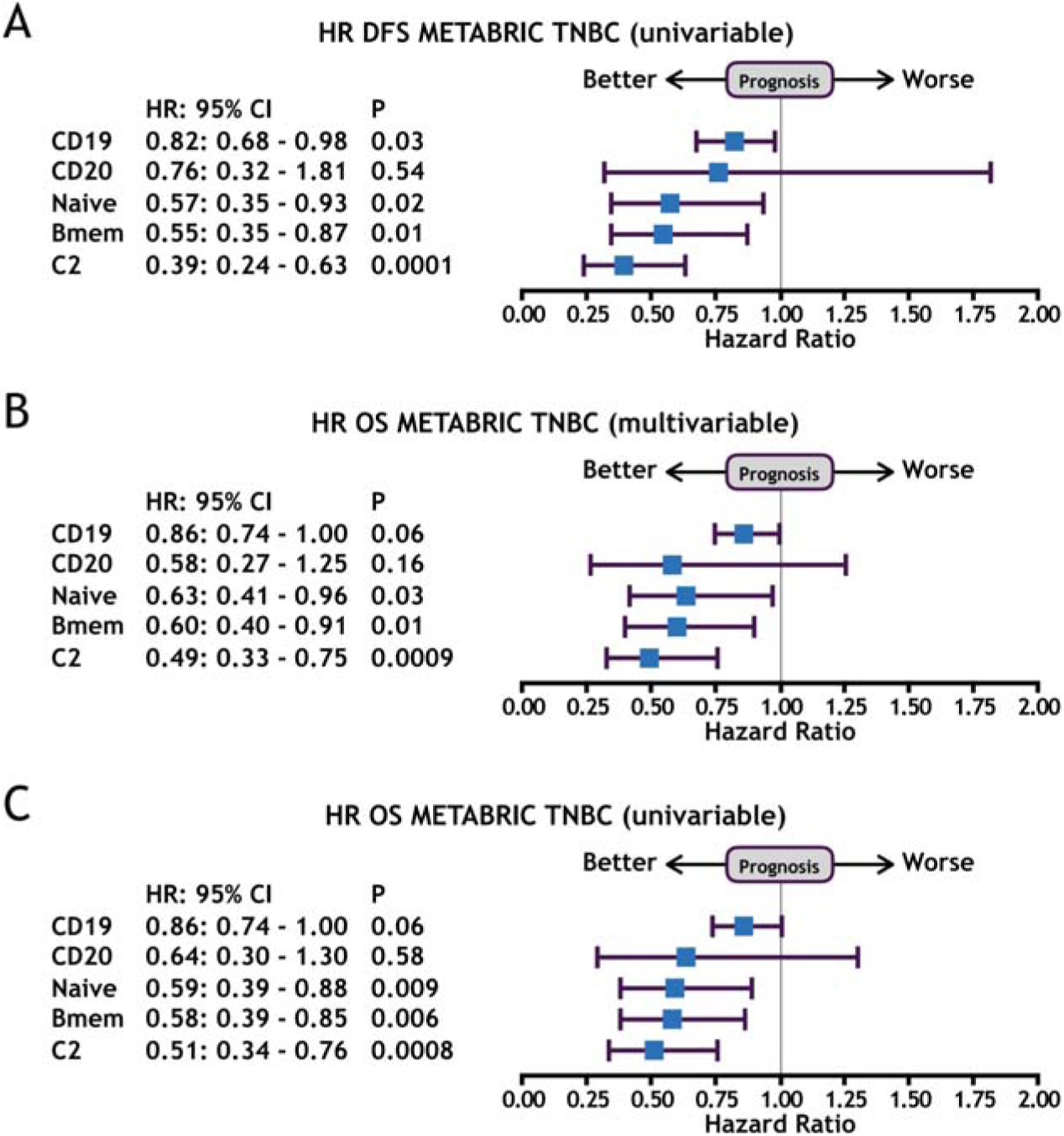
Prognostic effect of CD19, CD20, Naïve B, memory B and C2 Naïve B signatures in TNBC patients. Forest plots of hazard ratios for univariable disease free survival (A), multivariable overall survival (B) and univariable overall survival (C). Forest plots showed hazard ratios (blue squares) and confidence intervals (horizontal ranges) derived from Cox regression survival analyses. Multivariable analysis was adjusted for lymph nodes status, tumor size, age at diagnosis, and histological grade.

**Figure S10.**
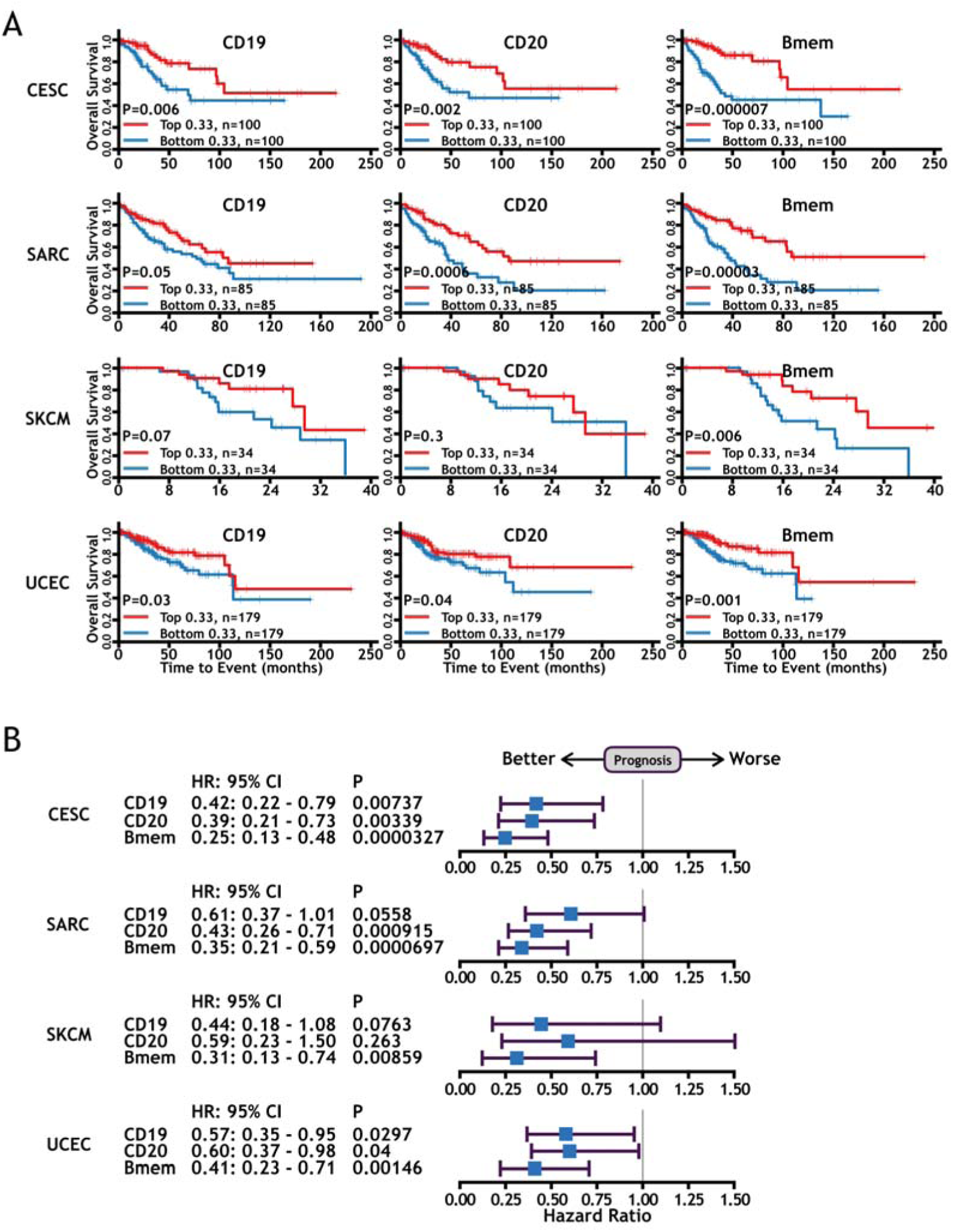
The memory B cell signature based on single-cell RNA-seq results also provide better prognostication than classic single B cell markers in other human cancer types. A. Kaplan–Meier survival curves for overall survival of CESC, SARC, SKCM, UCEC patients in TCGA project according to expression of CD19, CD20, and memory B signature. The p-values were calculated by log-rank test. B. Prognostic effect of CD19, CD20, and memory B signature in CESC, SARC, SKCM, UCEC patents. Forest plots show HRs (blue squares) and confidence intervals (horizontal ranges) derived from Cox regression survival analyses for overall survival in univariable analysis.

Table S1. **Statistic information of the single cell RNA-sequencing libraries**

Table S2. **Statistic information of the single cell BCR and TCR libraries**

Table S3. **Summary of whole-exome sequencing results**

Table S4. **Marker genes for 4 B cell clusters in low resolution**

Table S5. **Marker genes for 13 B cell clusters in high resolution**

Table S6. **Signature genes of Bmem, Naïve B and C2 Naïve B**

Table S7. **Survival analysis in TCGA samples**

## Author Contributions

Q.H., Y.H., P.Q., and Y.Z. conceived the study. Q.H., Y.H., P.Q., G.L., X.M., L.G., Z.J., J.W., and T.C. performed experiments and analyzed data. S.X., X.H., Y.G., and P.Q. collected patient samples. Y.Z. analyzed data and wrote the manuscript with support from all authors.

## Acknowledgments

We thank all breast cancer patients participated in this study at Xinxiang central hospital. We thank members of Y.Z. lab for helpful discussion and support. The research in Y.Z. lab is supported by National Natural Science Foundation of China (81572795, 81773304), the “Hundred, Thousand and Ten Thousand Talent Project” by Beijing municipal government (2015001, 2017A02), “National Thousand Young Talents Program” of China. Q.H. is supported by National Natural Science Foundation of China (31701135). P.Q. is supported by 2018 Joint Construction Project of Henan Medical Science and Technology Tackling Plan. We thank the municipal government of Beijing and the Ministry of Science and Technology of China for funds allocated to NIBS.

